# Molecular Predicting Drought Tolerance in Maize Inbred Lines by Machine Learning Approaches

**DOI:** 10.1101/578880

## Abstract

Drought is one of the prime abiotic stresses in the world. Now, amongst the new technologies available for speed up the releasing of new drought tolerance genotypes, there is an emanate discipline called machine learning. The study presents Machine Learning for identification, classification and prediction of drought tolerance maize inbred lines based on SSR genetic markers datasets generated from PCR reactions. A total of 356 SSR reproducible fragment alleles were detected across the 71 polymorphic SSR loci. A dataset of 12 inbred lines with these fragments prepared as attributes and was imported into RapidMiner software. After removal of duplicates, useless and correlated features, 311 feature attributes were polymorphic, ranging in size from 1500 to 3500 bp. The most important attribute fragment alleles in different attribute weighting selected. Ten datasets created using attribute selection (weighting) algorithms. Different classification algorithms were applied on datasets. These can be used to identify groups of alleles with similar patterns of expression, and are able to create some models that have been applied successfully in the prediction, classification and pattern recognition in drought stress. Some unsupervised models were able to differentiate tolerant inbred lines from susceptible. Four unsupervised models were able to produce the different decision trees with root and leaves. The most important attribute alleles almost in all of models were phi033a3, bnlg1347a1 and bnlg172a2 respectively, that can help to identify tolerant maize inbred lines with high precision.

## Introduction

Drought is the main cause of reduced yields in cereal crops (Edmeades et al., 2000), after low soil fertility, the second most important cause of yield loss for maize (Dariano et al., 2016). It is a major limitation in maize production, particularly under climatic changes (Lobell et al., 2014), leading to a grain yield reduction of 25–30%, even without any harvests during extremely severe drought events (Khazaei et al., 2013; Singh et al., 2016). Drought has greater effect despite cultivar and agronomic management improvements. Climatic warming is expected to further enhance the harmful impact of droughts; potentially leading to a significant decrease in maize yields (Ribaut et al., 2009; Lobell et al., 2014). Thus, research on maize drought tolerance and its mechanisms remain hotspot under the pressure of increasing environmental conditions that reflect the impact of human activities on the environment that leads to the appearance of environmental problems. (Lobell et al., 2014; Khazaei et al. 2013; Ribaut et al., 2009). Drought has even been thought of as a “cancer” to plants, owing to its complexity and destructiveness. Hence, there is enormous interest in and demand for improving maize drought tolerance through biotechnology (Wang et al., 2016).

Maize respond drought stress at the physiological, biochemical, and molecular levels to adapt to changing environmental conditions (Jaglo-Ottosen et al., 1998; Shinozaki et al., 2003; Xiong et al., 2002). The identification and use of molecular markers to assist in selection of quantitative trait loci, genome-wide selection, and association mapping have become common areas of research. In the future, the destructive impact of drought may grow, as the spectre of climate change becomes a reality. With the development of PCR-based DNA markers such as RAPD (Raibut 2009) SSR (Ercisli et al., 2011), AFLPs (Pafundo et al., 2005) and single nucleotide polymorphism (Reale et al. 2006), marker technology today offers a palette of powerful tools to analysis the plant genome. (Tardieu 2012; Wang et al. 2011; Yadav et al., 2011). They have enabled the identification of genes and genome associated with the expression of qualitative and quantitative traits and has led to a better understanding of the complex genome of various plants (Shinozaki and Yamaguchi-Shinozaki, 2007; Tao et al., 2011), besides helping in identifying the desired species at any growth stage of the plant. Despite numerous published reports of molecular markers for drought-related traits, practical applications of such results in maize improvement are scarce (Benesova et al., 2012) and results are not completely satisfying and more research on methodologies is needed (Ornella et al., 2012).

Machine learning is an emerging inherently multidisciplinary approach to data analysis with a revolutionary impact on a variety of areas and refers to a group of computerized modelling approaches that can learn patterns from the data to make automatic decisions without programming obvious rules. (Dror et al., 2005; Hajiloo et al., 2013; Hepworth et al., 2012, Hor et al., 2013; Mutka and Bart, 2015; Smith et al., 2013; Tran, 2014; Yan et al., 2013). The main idea of machine learning is to effectively utilize experiences to discover main underlying structures, similarities, or dissimilarities present in data to explain or classify a new experience properly. A key ability of machine learning tools, in its most fundamental form, is their ability to generalize complex patterns and making intelligent decisions from data. (Sing et al., 2016; Forghani and Yazdi, 2014; Ma et al., 2012). Machine learning will also enhance our understanding of pathogen–plant interactions as well as the interaction of plants with other stresses (Kuska et al. 2015; Sing et al. 2016; Romer et al., 2012). One of the major advantages of machine-learning approaches for plant breeders and biologists is the opportunity to search datasets to discover patterns and govern discovery by simultaneously looking at a combination of factors instead of analysing each feature individually (Mutka and Bart, 2015; Sing et al., 2016).

Due to their high generalization capabilities and distribution-free properties they are presented as a valuable alternative to traditional statistical techniques applied in maize breeding, even the more recently introduced linear mixed models (Maenhout et al., 2010; Maenhout et al., 2007; Ornella, Cervigni and Tapia, 2012).

In this study, using polymorphic SSR loci, molecular analysis of the selected set of inbred lines carried out. After detecting SSR reproducible fragment alleles, datasets were created with different attribute weighting methods. Different unsupervised and supervised classification algorithms run on datasets. Machine learning models differentiate tolerant maize inbred lines from susceptible. The main objective of this study is to develop machine-learning models in the field of plant stress phenotyping in identification, classification, quantification and prediction. Understanding the genetic basis of drought tolerance of maize with major attribute alleles may facilitate efforts to improve drought tolerant inbred lines.

## Materials and Methods

A selected set of 12 inbred lines with distinct tolerance and susceptibility responses to drought stress at the flowering stage in maize were analyzed in this study. These include four DTPY lines (DTPY65, DTPY108, DTPY116 and DTPY194), three DTPW lines (DTPW51, DTPW105, DTPW183 and DTPW195) besides Ac7643, Ac7729, CM139 and CM140. Genomic DNA from each of genotypes was extracted from a bulk of 10-15 plants, using a modified CTAB method (Saghi-Maroof et al., 1994) with suitable modifications. Primers for SSR loci were custom synthesized through Research Genetics Inc., USA or Sigma Aldrich, UK, based on forward and reverse primer sequence information available in public domain (MaizeDB; http://www.agron.missouri.edu).

Molecular analysis of the selected set of inbred lines was carried out using 71 polymorphic SSR loci covering various bin (chromosome) locations (Table 1). In addition, SSR markers located at specific bin locations with potential utility for QTL mapping of drought tolerance in maize were utilized in the study (Table 1). PCR reactions were carried out in 10 µl reaction volume containing of 10× PCR buffer (10 mM Tris-HCl pH 8.3, 50 mM KCl), 1.5 mM MgCl_2_, 0.2 mM each dNTPs, 5 pmol of each forward and reverse primer, 0.5U of Taq DNA polymerase (NEB), and 5 ng of genomic DNA. Reactions were carried out in GenAmp PCR system 9700 (Applied Biosystems, USA) thermal cycler using the following temperature profile: an initial denaturation of 5 min at 94°C followed by 35 cycles of 45 s at 94°C, 45 s at 55°C and 1 min at 72°C, then a final extension of 5 min at 72°C. Amplification products were resolved on 3% metaphor agarose gels using a horizontal gel electrophoresis unit (CBS Scientific, USA). The DNA fragments were then visualized under UV-transilluminator and documented using ALPHA IMAGER gel documentation system (Alpha Innotech, USA) which was stored for further scoring and permanent records. The fragment alleles were visually scored as present (1) or absent (0).

**Table 1.** Name, repeat type and bin location of SSR marker

| SSR Locus | Repeat Type | Bin Location | SSR Locus | Repeat Type | Bin Location | SSR Locus | Repeat Type | Bin Location |
| --- | --- | --- | --- | --- | --- | --- | --- | --- |
| phi097 | TAG | 1.01 | bnlg420 | CT | 3.05 | phi089 | ATGC | 6.08 |
| bnlg147 | CA | 1.02 | phi088 | ACT | 3.08 | bnlg339 | CT | 7.03 |
| bnlg1016 | AG | 1.04 | bnlg1182 | AG | 3.09 | bnlg155 | CT | 7.04 |
| bnlg1556 | AG | 1.06 | bnlg2244 | AG | 4.03 | phi116 | AGTG-ACG | 7.06 |
| phi037 | AG | 1.08 | bnlg1729 | AG | 4.05 | phi119 | AG | 8.02 |
| bnlg400 | AG | 1.09 | bnlg252 | AG | 4.06 | phi014 | GGC | 8.04 |
| bnlg1347 | AG | 1.1 | bnlg589 | AG | 4.11 | bnlg162 | CA | 8.05 |
| bnlg504 | CA | 1.11 | phi113 | GTCT | 5.02 | bnlg240 | AG | 8.06 |
| umc1331 | GGT | 1.11 | bnlg2323 | AG | 5.04 | bnlg1065 | AG | 8.06 |
| bnlg1092 | AG | 2.01 | phi101 | ACT | 5.06 | phi015 | AAAC | 8.08-8.09 |
| bnlg2042 | AG | 2.02 | bnlg389 | CT | 5.09 | phi080 | AGGAG | 8.08-8.09 |
| bnlg1297 | AG | 2.02 | bnlg1043 | AG | 6 | bnlg1272 | AG | 9 |
| bnlg1064 | AG | 2.03 | phi077 | AG | 6.01 | phi033 | AAG | 9.01 |
| umc1555 | TTCA | 2.03 | umc1018 | CT | 6.01 | bnlg1730 | AG | 9.03 |
| bnlg381 | CA | 2.03 | umc1006 | GA | 6.02 | bnlg1209 | AG | 9.05 |
| bnlg1613 | AG | 2.04 | phi070 | AGCTG | 6.03 | phi041 | AGCC | 10 |
| bnlg1175 | AG | 2.04 | umc1572 | GA | 6.03 | phi059 | ACC | 10.02 |
| dup21 | AG | 2.05 | umc1014 | GA | 6.04 | bnlg210 | AG | 10.03 |
| mage05 | AG | 2.05 | bnlg1154 | AG | 6.05 | bnlg1655 | AG | 10.03 |
| **bnlg121** | CT | 2.06 | **bnlg1702** | AG | 6.05 | **bnlg1518** | AG | 10.04 |
| **bnlg1138** | AG | 2.06 | **mmc0241** | TA-N-TG | 6.05 | **bnlg1526** | AG | 10.04 |
| **bnlg1045** | AG | 2.07 | **phi078** | AAAG | 6.05 | **bnlg594** | AG | 10.06 |
| **bnlg198** | CT | 2.08 | **phi102** | AT | 6.05 | **phi035** | AC | 10.06 |
| **bnlg1520** | AG | 2.09 | **nc013** | AG | 6.05 |  |  |  |

Allele designations were made for the amplification products for each SSR locus was determined based on the positions of the bands relative to the 100 bp-molecular-weight ladder (Fig. 1). A total of 356 SSR reproducible fragment alleles were detected across the 71 polymorphic SSR loci. A dataset of 12 inbred lines with these fragments prepared as attributes and was imported into RapidMiner software [RapidMiner 5.2, Rapid-I GmbH, Stochumer Str. 475, 44227 Dortmund, Germany]. Then, the steps detailed below were applied to this dataset.

**Figure 1.**
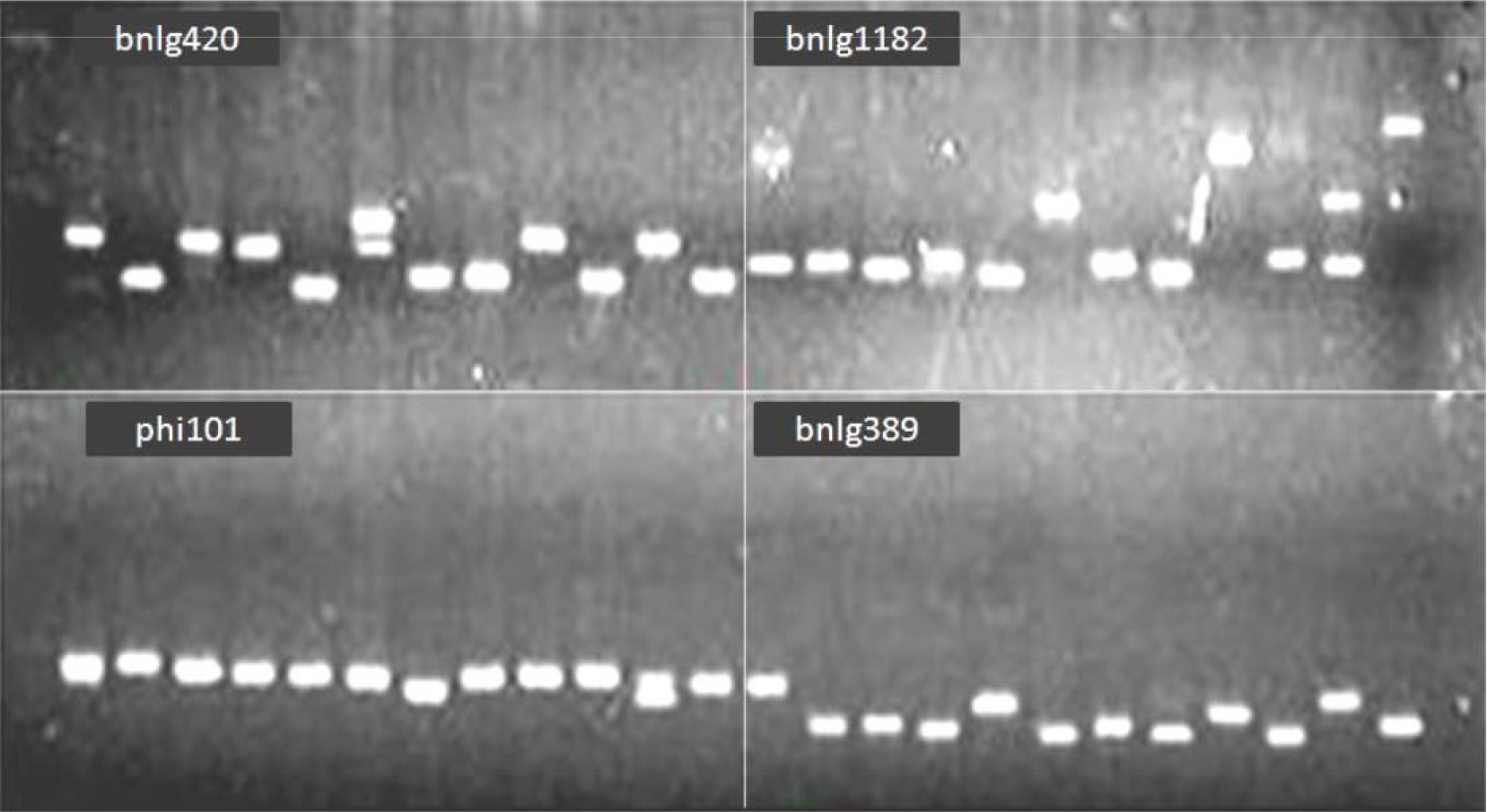
SSR fragment alleles can often be distinguished on agarose gels.

### Data cleaning

Useless attributes were removed from the dataset. Nominal attributes were regarded as useless when the most frequent values were contained in more or less than nominal useless above or below per cent of all examples. After cleaning, this database was labelled the final cleaned database (FCdb).

### Attribute Weighting

To identify the most important allele attributes and to find the possible patterns in features that contribute to different maize inbred lines, 10 different algorithms of attribute weightings including *Information gain, Information Gain ratio, Rule*, Deviation, *Chi squared statistic, Gini index, Uncertainty, Relief, SVM and PCA* were applied to the FCdb. *Weight by Information gain* operator calculated the relevance of a feature by computing the information gain in class distribution. *Weight by Information Gain ratio* calculated the relevance of a feature by computing the information gain ratio for the class distribution. *Weight by Rule* calculated the relevance of a feature by computing the error rate of a OneR Model on the example set without this feature. *Weight Deviation* operator created weights from the standard deviations of all attributes. The values were normalised by the average, the minimum, or the maximum of the attribute. *Weight by Chi squared statistic* operator calculated the relevance of a feature by computing, for each attribute of the input example set, the value of the chi-squared statistic with respect to the class attribute. *Weight by Gini index* operator calculated the relevance of an attribute by computing the Gini index of the class distribution, if the given example set would have been split according to the feature. *Weight by Uncertainty* operator calculated the relevance of an attribute by measuring the symmetrical uncertainty with respect to the class. *Weight by Relief* measured the relevance of features by sampling examples and comparing the value of the current feature for the nearest example of the same and of a different class. This version also worked for multiple classes and regression data sets. The resulting weights were normalised into the interval between 0 and 1. *Weight by SVM* operator used the coefficients of the normal vector of a linear SVM as feature weights. *Weight by PCA* operator used the factors of the first of the principal components as feature weights.

### Attribute selection

After attribute weighting models were run on the dataset, each alleles attribute (feature) gained a value between 0 and 1, which revealed the importance of that attribute with regards to a target attribute (susceptible-tolerance cultivar). All variables with weights higher than 0.50 were selected and 10 new datasets created and were named according to their attribute weighting models (*Information gain, Information gain ratio, Rule, Deviation, Chi Squared, Gini index, Uncertainty, Relief, SVM and PCA*. These newly formed datasets were used to join with subsequent models (supervised and unsupervised). Each model of supervised or unsupervised clustering were performed 11 times; the first time it ran on the main dataset (FCdb) and then on the 10 newly formed datasets from attribute weighting and selection.

### Unsupervised clustering algorithms

Ten newly created datasets generated as the outcomes of different attribute weighing algorithms along with FCdb were applied to K-Means, K-Medoids, *Support Vector Clustering (SVC)* and *Expectation Maximization (EM)* clustering algorithms. *K-Means* uses kernels to estimate the distance between objects and clusters. Because of the nature of kernels, it is necessary to sum over all elements of a cluster to calculate one distance. *K-Medoids* represents an implementation of k-Medoids. This operator will create a cluster attribute if it is not yet present. SVC represents implementations of Support Vector algorithm which will create a cluster attribute if not present yet. *EM* represents an implementation of the EM-algorithm.

### Supervised Classification

Three classes of supervised classification (*Decision Trees*, SVM and *Baysian* models) applied as follows. To calculate the accuracy of each model, 10-fold cross validation is used to train and test models on all patterns. To perform cross validation, all the records were randomly divided into five parts; four sets were used for training and the 5th one for testing. The process was repeated five times and the accuracy for true, false and total accuracy calculated. The final accuracy is the average of the accuracy in all five tests.

### Decision Trees

Six tree induction models including *Decision Tree, Decision Tree Parallel, Decision Stump, Random Tree, ID3 Numerical and Random Forest* were run on the main dataset (FCdb). Each tree induction model ran with the following four different criteria: *Gain Ratio, Information Gain, Gini Index and Accuracy*. In addition, a *weight-based parallel decision tree* model, which learns a pruned decision tree based on an arbitrary feature relevance test (attribute weighting scheme as inner operator), was run with 13 different weighing criteria (*SVM, Gini Index, Uncertainty, PCA, Chi Squared, Rule, Relief, Information Gain, Information Gain Ratio, Deviation, Correlation, Value Average, and Tree Importance*). The accuracy of each tree computed based on the previous explanation.

### Support Vector Machine Approach

Support Vector Machines (SVMs) are popular and powerful techniques for supervised data classification and prediction; so SVM, LibSVM, SVM Linear and SVME used here to implement different models to predict maize cultivars based on Susceptible-Tolerance features. Briefly, main database (FCdb) transformed to SVM format and scaled by grid search (to avoid attributes in greater numeric ranges dominating those in smaller numeric ranges) and to find the optimal values for operator parameters. To prevent overfitting problems, 5-fold cross validation applied. Dataset divided into 5 parts and 4 parts used as training set and the last part as testing set. The procedure repeated for 10 different testing sets and the average of accuracy computed. RBF kernel that nonlinearly maps samples into a higher dimensional space and can handle the case when the relation between class labels and attributes is nonlinear used to run the model. Other kernels such as linear, poly, sigmoid and pre-computed were also applied to the dataset to find the best accuracy.

### Naïve Bayes

*Naïve Bayes* based on *Bayes* conditional probability rule was used for performing classification tasks (Wang and Tseng 2013). *Naïve Bayes* assumes the predictors are statistically independent which makes it an effective classification tool that is easy to interpret. Two models, *Naïve base* (returns classification model using estimated normal distributions) and *Naïve base kernel* (returns classification model using estimated kernel densities) used and the model accuracy in predicting the right resistance - susceptible computed as stated before.

## Results

SSR alleles differing by several repeat units can often be distinguished on agarose gels. High level of allelic variation, making SSR valuable as genetic attributes (Fig. 1). SSR is used for understanding of the evolutionary genetics and sequences controlling traits of economic interest of maize (Xu et al., 2009).

As mentioned in Materials and Methods, the initial dataset contained 12 maize inbred lines with 356 SSR fragment attributes. Following removal of duplicates, useless attributes, and correlated features (data cleaning) 311 features remained, meaning these fragment attributes were polymorphic, ranging in size from 1500 to 3500 bp.

### Attribute weighting

The number of attributes gained weights higher than 0.5 in each weighting model were as follows: *Relief* 168, Rule 168, Deviation 147, SVM 134, *PCA* 93, *Info gain ratio* 20, *Uncertainty* 20, *Chi squared* 9, *Gini index* 8 and *Info Gain* 8 (Table 2). The most important attribute fragment allele in all of models was phi033a3, which weighted equal to 1.0 in all weighting models except to PCA and can help to identify tolerance maize inbred lines from susceptible.

**Table 2.**
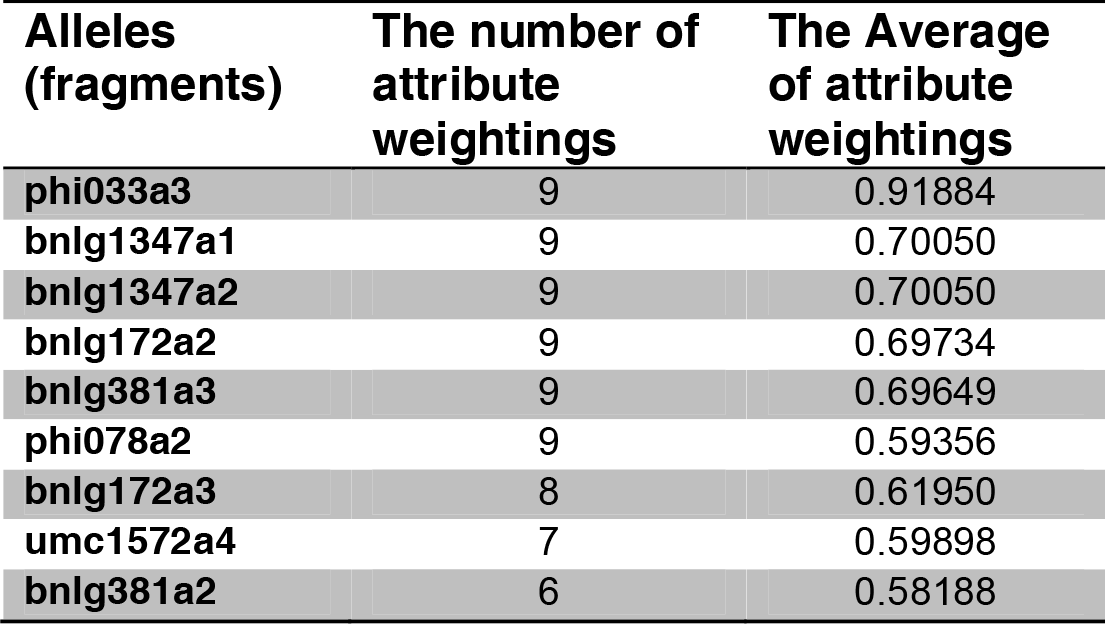
The number and the average of most important alleles (fragments) selected by different attribute weighting algorithms.

| **Alleles**<br>**(fragments)** | **The number of**<br>**attribute**<br>**weightings** | **The Average**<br>**of attribute**<br>**weightings** |
| --- | --- | --- |
| **phi033a3** | 9 | 0.91884 |
| **bnlg1347a1** | 9 | 0.70050 |
| **bnlg1347a2** | 9 | 0.70050 |
| **bnlg172a2** | 9 | 0.69734 |
| **bnlg381a3** | 9 | 0.69649 |
| **phi078a2** | 9 | 0.59356 |
| **bnlg172a3** | 8 | 0.61950 |
| **umc1572a4** | 7 | 0.59898 |
| **bnlg381a2** | 6 | 0.58188 |

In weighting by *Deviation* operator 46 attribute alleles including bnlg2323a1, bnlg1730a4, phi088a3, bnlg1655a2, mage05a3 and bnlg210a3 weighed equal to 1.0. In Weighting by *Rule* model five attributes, phi033a3, bnlg172a3, phi078a2, bnlg381a3 and bnlg172a2; in *SVM* model three attributes, phi033a3, bnlg1347a1 and bnlg1347a2; and in *PCA model* five attributes, bnlg1138a1, bnlg18a3, phi102a3 and phi102a2 weighted equal to 1.0. These attributes were the most important attributes selected when these models applied on dataset of correlated removed features. Five of ten attribute weighting models including Uncertainty, Info Gain Ratio, Gini Index, Information Gain and Chi Squared selected the following attribute alleles: phi033a3, bnlg172a2, bnlg1347a1, bnlg1347a2, umc1572a4, bnlg172a3 and bnlg381a3 weighted more than 0.7. In Weighting by *Releif* model only phi033a3 attribute allele weighted equal to 1.0. These attributes were the most important attributes selected when these models applied on dataset of correlated removed features (Table3). Almost all of these attributes are the main branches of decision trees (Fig. 3 and Fig. 4).

**Table 3.** The attribute weighting models and the numbers of important fragment alleles selected by each model and the most important variables selected by each attribute weighting algorithms.

| Attribute Weighting | Number of Variable | Important Attribute Fragment Alleles |  |  |  |  |  |
| --- | --- | --- | --- | --- | --- | --- | --- |
| **Rule** | 168 | phi033a3 | bnlg172a3 | phi078a2 | bnlg381a3 | bnlg172a2 | umc1572a4 |
| **Deviation** | 147 | bnlg2323a1 | bnlg1730a4 | phi088a3 | bnlg1655a2 | mage05a3 | bnlg210a3 |
| **SVM** | 133 | phi033a3 | bnlg1347a1 | bnlg1347a2 | bnlg172a2 | bnlg381a3 | bnlg1154a3 |
| **PCA** | 92 | bnlg1138a1 | phi035a3 | bnlg18a3 | phi102a3 | phi102a2 | bnlg381a3 |
| **Uncertainty** | 20 | phi033a3 | bnlg172a2 | bnlg1347a1 | bnlg1347a2 | umc1572a4 | bnlg172a3 |
| **InfoGainRatio** | 20 | phi033a3 | bnlg172a2 | bnlg1347a1 | bnlg1347a2 | umc1572a4 | bnlg172a3 |
| **Gini Index** | 8 | phi033a3 | bnlg172a2 | bnlg1347a1 | bnlg1347a2 | umc1572a4 | bnlg172a3 |
| **Relief** | 8 | phi033a3 | bnlg172a2 | bnlg1347a1 | bnlg1347a2 | dup21a4 | bnlg252a2 |
| **Info Gain** | 8 | phi033a3 | bnlg172a2 | bnlg1347a1 | bnlg1347a2 | umc1572a4 | bnlg172a3 |
| **Chi Squared** | 7 | phi033a3 | bnlg172a2 | bnlg1347a1 | bnlg1347a2 | umc1572a4 | bnlg172a3 |

### Unsupervised Clustering Algorithms

Three different unsupervised clustering algorithms (*K-Means, K-Medoids and SVC*) were applied on ten datasets created using attribute selection (weighting) algorithms. Some models, such as the application of the *SVC* algorithm on all datasets were able to differentiate **tolerant** inbred lines from susceptible. In this algorithm all tolerant inbred lines predicted as correct class of tolerance (Fig. 2-A). Application of the *K-Means* to *SVM*, *PCA* and *Deviation* databases was able to assign respectively 66.7%, 33.3% and 50% drought tolerance inbred lines into its correct class while application of *K- Medoids* to same database was able to assign respectively 50%, 83.3% (Fig. 2-C) and 100% (Fig. 2-B) drought tolerance inbred lines into its correct class (Table 4). Combination of *K-Medoids* clustering method with *Deviation* attribute weighting selected the right classes of drought tolerance inbred lines (Fig. 2-B) while Combination of *K-Means* clustering method with *PCA* attribute weighting selected the right classes of drought susceptible inbred lines (Fig. 2-D).

**Figure 2.**
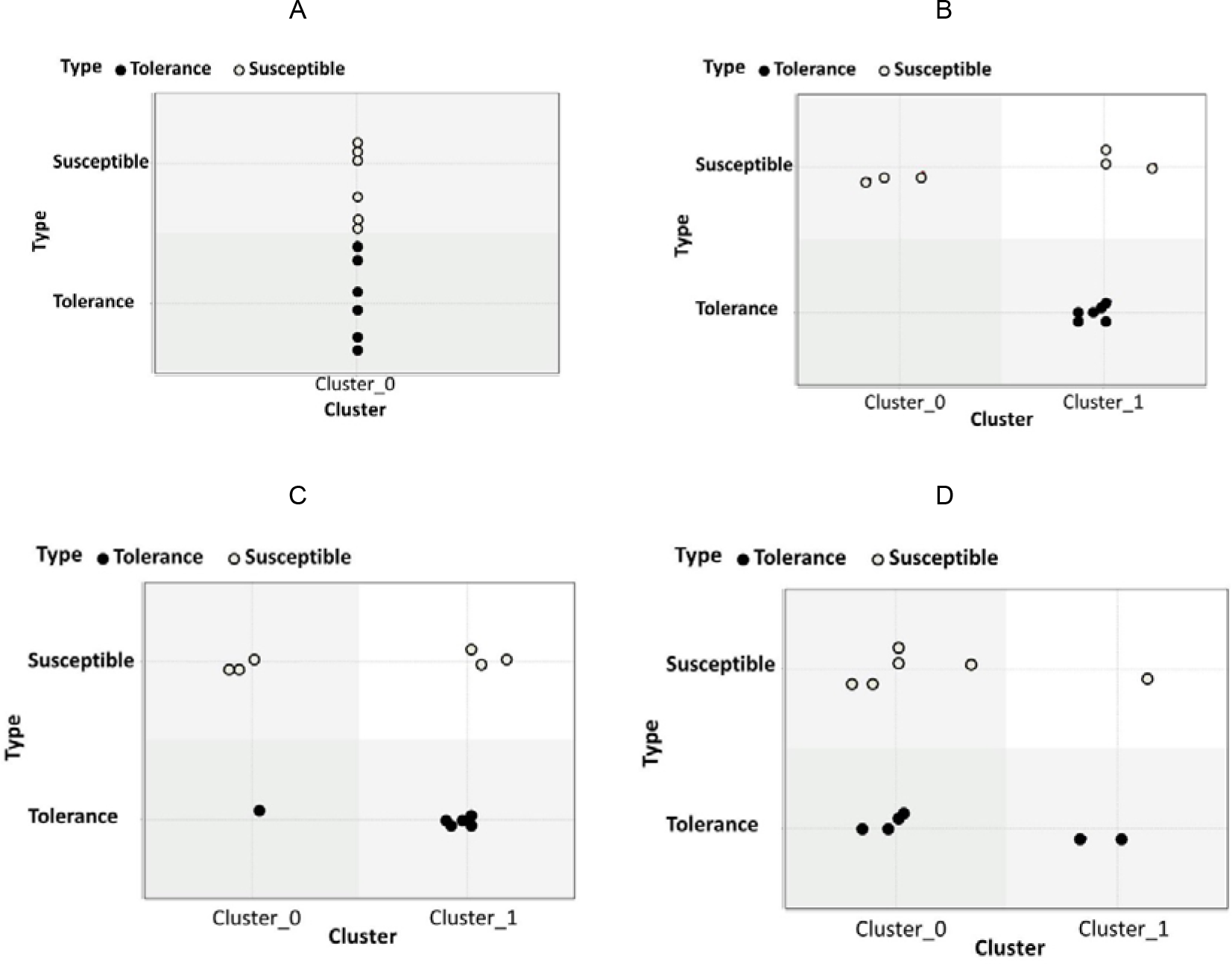
Application of the SVC algorithm on ten datasets (A) was unable to categorize inbred lines into correct clusters; K-Medoids algorithm to the Deviation (B) and to PCA(C) was able to categorize drought tolerance inbred lines into correct clusters and K-means algorithm to the PCA (C) was able to categorize susceptible tolerance inbred lines into correct clusters.

**Table 4.** Percent correct class clustering of the K-Means and K-Medoids to SVM, PCA and Deviation databases

| Unsupervised Algorithm | Dataset | Susceptible | Tolerance |
| --- | --- | --- | --- |
| **K-Means** | SVM | 33.3 | 66.7 |
|  | PCA | 83.3 | 33.3 |
|  | Deviation | 66.7 | 50.0 |
| **K-Medoids** | SVM | 66.7 | 50.0 |
|  | PCA | 50.0 | 83.3 |
|  | Deviation | 50.0 | 100.0 |

### Supervised Classification

#### Decision Trees

From six induction models *Random Tree* and *ID3Numerical* were not able to produce decision trees. *Decision Stump*, *Decision Tree*, *Decision Parallel* and *Random Forest Tree* with 4 from 13 different criteria; *gain ratio*, *information gain*, *gini index* and *accuracy* (run on 11 datasets) were able to produce the different decision trees with root and leaves (Figure 3 - Figure 5). *Decision Stump*, *Decision Tree* and *Decision Parallel* in 4 criteria *Gain Ratio*, *Information Gain*, *Gini Index* and *Accuracy* almost on all databases can produce the same simple decision tree (Figure 3), but their overall accuracy is below 60%. In this decision tree phi033a3 attribute fragment allele with the average attribute weighting of 0.92 was the sole fragment allele used to build this tree. If there is phi 033a3 allele, the maize inbred line is tolerance to drought; otherwise, the inbred line is susceptible.

**Figure 3.**
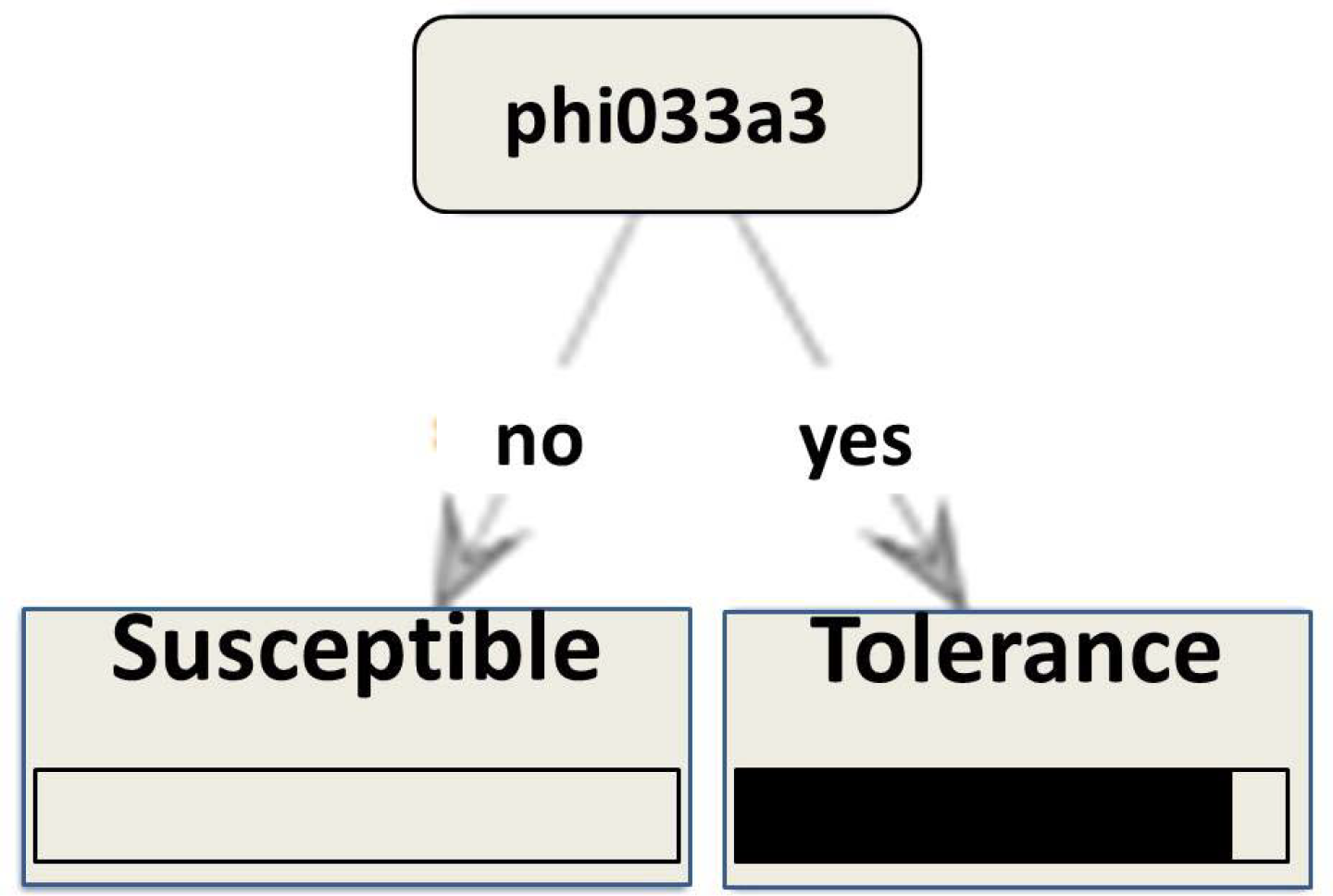
Almost Decision Stump, Decision Tree and Decision Parallel in 4 criteria gain ratio, information gain, gini index and accuracy on all databases can produce the same simple decision tree.

*Random Forest* model with four criteria on *Chi Square*, *Deviation*, *Information Gain* and *Gini Index* datasets were able to produce different decision trees with overall accuracy above 50% (Table5). Decision trees of *Random Forest* model with *Accuracy* criterion run on *Information Gain* dataset (Fig. 4 -A) and *Gini Index* Criterion run on *Gini Index* dataset (Fig. 4 -B) show 100% overall accuracy and precision. Figure 4 -A shows that bnlg1347a2 attribute fragment allele was the sole fragment allele used to build this tree. If there is bnlg1347a2 allele, the maize inbred line is drought tolerance; otherwise, the inbred line is susceptible. Figure 4 -B shows that bnlg381a3 fragment allele was the sole fragment allele used to build this tree. If there is bnlg381a3 allele, the maize inbred line is tolerance. *Random Forest* model with *Chi Square* dataset was able to produce different decision trees with overall accuracy 90% and overall precision 100% when run with *Accuracy* (Fig. 5 -A), *Gain Ratio* (Fig. 5 -B) and *Gini Index* criteria (Fig. 5 -C). *Random Forest* model with *Gini Index* criterion also produce decision tree (Fig. 5 -D) with overall accuracy 90% and overall precision 100% when run on *Information Gain* dataset.

**Figure 4.**
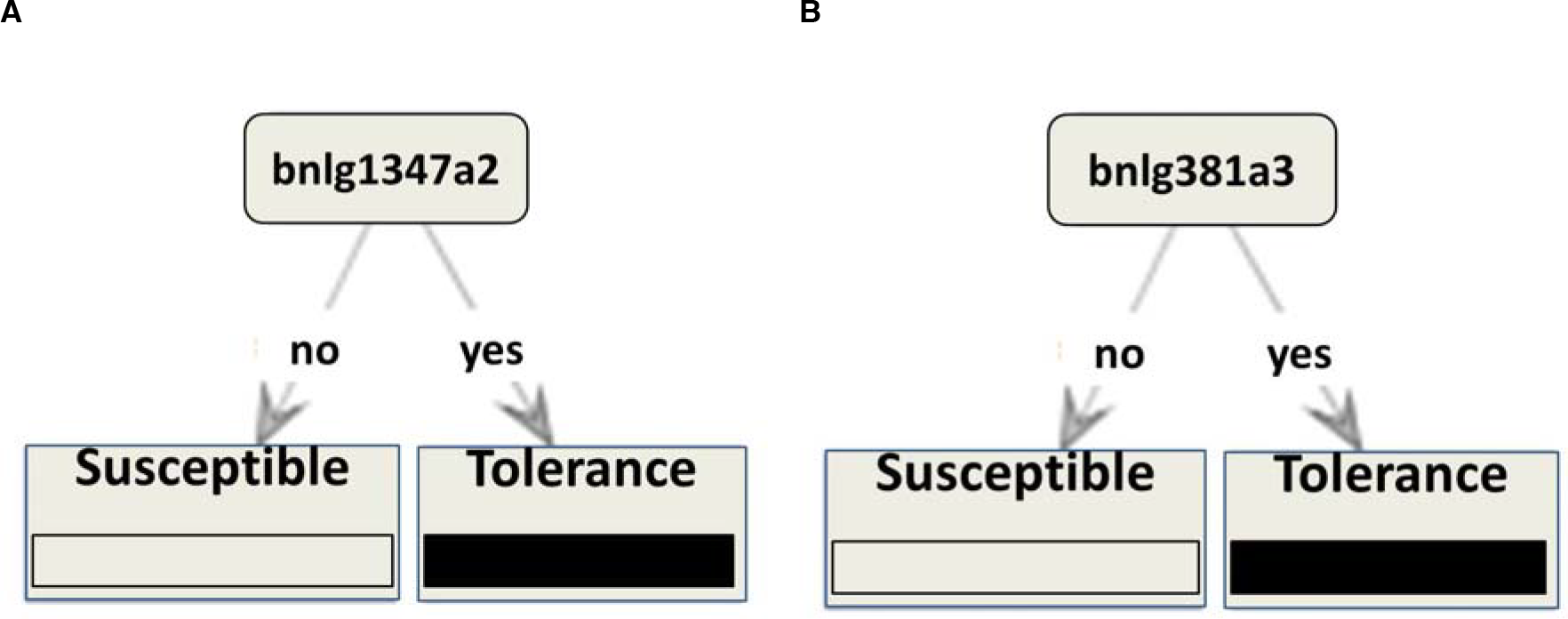
Decision trees on Random Forest model with Accuracy criterion run on Information Gain dataset (A) and Gini Index Criterion run on Gini Index dataset (B).

**Table 5.** Overall Accuracy and Precision; Precision and Recall Tolerance and Susceptible in Random forest with 4 criteria run on Chi Squre, Deviation, Information Gain and Gini Index Datasets in decision trees

| Dataset | Criteria | Precision Tolerance | Precision Susceptible | Recall Tolerance | Recall Susceptible | Overall Accuracy | Overall Precision |
| --- | --- | --- | --- | --- | --- | --- | --- |
| **Chi Squire** | Accuracy | 86.0 | 100.0 | 100.0 | 83.3 | 90.0 | 100.0 |
|  | Gain ratio | 86.0 | 100.0 | 100.0 | 83.3 | 90.0 | 100.0 |
|  | Gini Index | 86.0 | 100.0 | 100.0 | 83.3 | 90.0 | 100.0 |
|  | Info. Gain | 75.0 | 100.0 | 100.0 | 66.7 | 85.0 | 100.0 |
| **Deviation** | Accuracy | 57.0 | 60.0 | 67.0 | 50.0 | 65.0 | 60.0 |
|  | Gain ratio | 50.0 | 50.0 | 67.0 | 33.0 | 45.0 | 50.0 |
|  | Gini Index | 80.0 | 71.0 | 67.0 | 83.0 | 75.0 | 71.0 |
|  | Info. Gain | 33.0 | 0.00 | 50.0 | 0.00 | 25.0 | 0.00 |
| **Info. Gain** | Accuracy | 100 | 100.0 | 100 | 100.0 | 100.0 | 100.0 |
|  | Gain ratio | 75.0 | 100.0 | 100.0 | 67.0 | 80.0 | 100.0 |
|  | Gini Index | 86.0 | 100.0 | 100.0 | 83.0 | 90.0 | 100.0 |
|  | Info. Gain | 75.0 | 100.0 | 100.0 | 67.0 | 85.0 | 100.0 |
| **Gini Index** | Accuracy | 75.0 | 100.0 | 100.0 | 67.0 | 80.0 | 100.0 |
|  | Gain ratio | 67.0 | 100.0 | 100.0 | 50.0 | 70.0 | 100.0 |
|  | Gini Index | 100.0 | 100.0 | 100.0 | 100.0 | 100.0 | 100.0 |
|  | Info. Gain | 71.0 | 80.0 | 83.0 | 67.0 | 70.0 | 80.0 |

### SVM approach

The result of the *SVM* prediction system based on the 10 fold cross validation sets show that overall accuracies were in the range of 0.0% to 100 %. The overall accuracies and prediction, prediction susceptible and tolerance precision and true susceptible and tolerance recall of 4 algorithms on *Chi Square*, *Gini Index*, *Information Gain*, *Relief* and *Uncertainty* databases and *LibSVM* algorithm on *Uncertainty* dataset were 100% (Table 6). The number of support vectors was 10-12 and for both susceptible and tolerance cultivars was 5-6 in these algorithm. These results demonstrated the manifest improvement of the prediction accuracies due to the application of generated datasets with *Support Vector Machine* model.

**Table 6.**
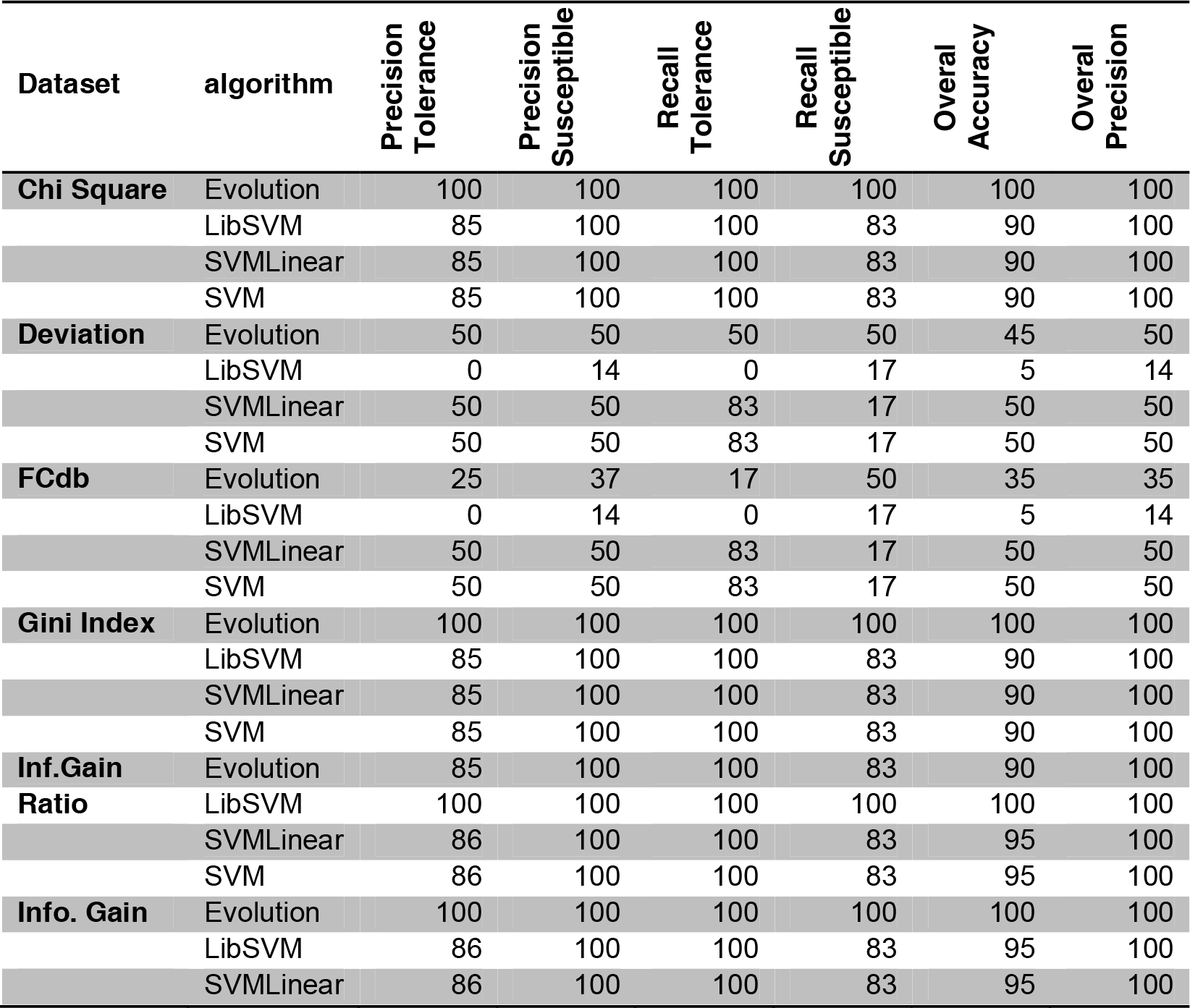
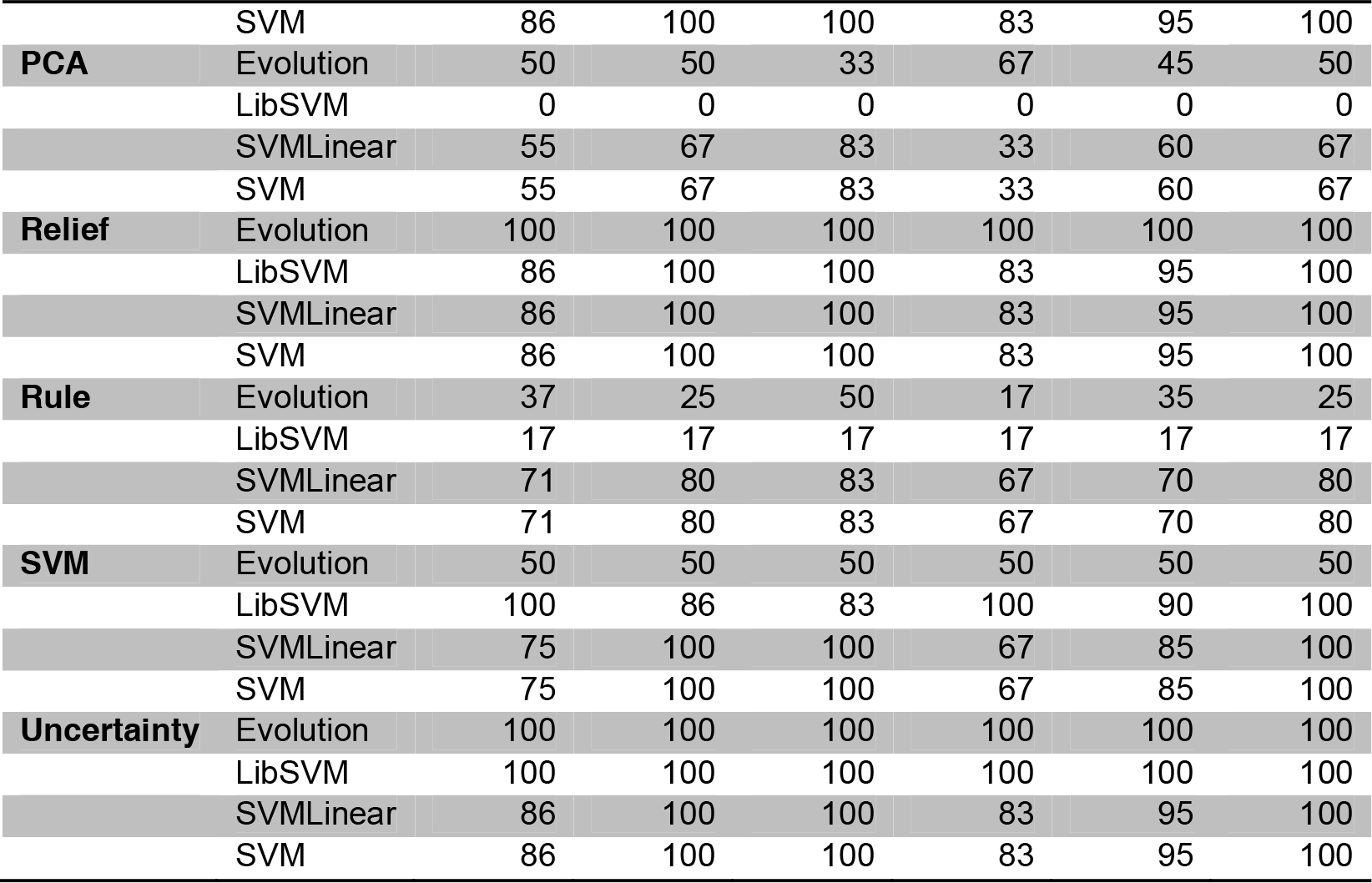
The accuracy, precision and recall of different models of SVM with different databases

### Naïve Bayes

The overall accuracies and prediction, prediction susceptible and tolerance precision and true susceptible and tolerance recall of *Naïve Base* and *Naïve Kernel* run on all databases were 100% except *Deviation*, *FCdb*, *PCA* and *Rule* datasets (Table 7). These results show the manifest improvement of the prediction accuracies due to the application of generated datasets with *Naïve Base* and *Naïve Kernel* models. Current practices rely heavily on the classical *Naïve Bayes* algorithm due to its simplicity and robustness. However, results from these algorithms are satisfactory.

**Table 7.**
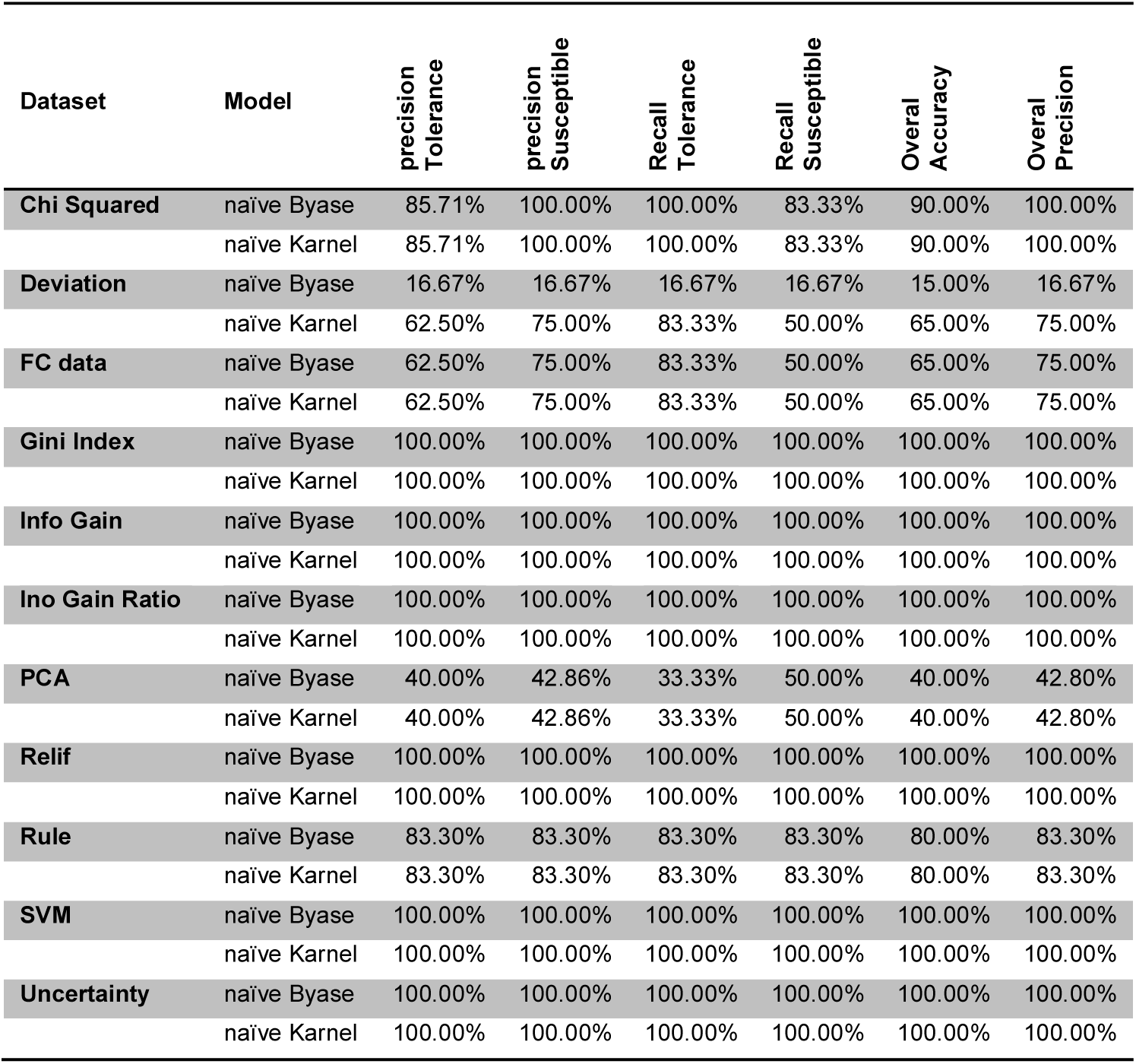
The accuracy, precision and recall of Naïve Bayse and Naïve kernel Models with different databases

## Discussion

The worldwide production of maize (*Zea mays* L.) is frequently impacted by water scarcity and as a result, increased drought tolerance is a priority target in maize breeding programs (Liu et al., 2013). Understanding the response of a crop to drought is the first step in the breeding of tolerant genotypes (Benesova et al., 2012). It is clear that one of the main objectives of plant breeding is the development of genotypes adapted to stressful environments. Drought resistance is labelled as a ‘complex trait’ mainly when viewed by molecular biologists (Fleury et al. 2010; Prasanna et al. 2009; Ravi et al. 2011). Recent advances in genomics have provided powerful new tools for molecular dissection of drought tolerance in maize during last decades. This platform is capable of expressing hundreds and thousands of drought-responsive genes, which are up or down regulated under dehydration stress according to growth stage, plant organ or even time of day (Blum, 2011). Identify the function of the candidate genes towards drought resistance is difficult. Inter-disciplinary scientists have been trying to understand and dissect the mechanisms of plant tolerance to drought stress using a variety of approaches (Mir et al., 2012). In order to achieve useful results, researchers require methods that consolidate, store and query combinations of structured and unstructured data sets efficiently and effectively. Herein, we combined molecular biology with biological knowledge discovery to determine the most important features contribute to the clustering, classification and prediction of drought tolerance inbred lines from susceptible based on SSR data.

The first and most important steps in any data processing task is to verify that your data values are correct or, at the very least, conform to some a set of rules. Data cleaning deals with detecting and removing errors and inconsistencies from data and were used to remove redundancy and co-linearity, useless or duplicated attributes in order to improve the quality of data which results in a smaller database (Ashrafi et al., 2011; Beiki et al., 2012). More than 7% of the attribute alleles discarded when these algorithms were applied on the original dataset. Each attribute weighting system uses a specific pattern to define the most important SSR fragment alleles. Thus, the results may be different (Baumgartner et al., 2010), as has been highlighted in previous studies (Ashrafi et al., 2011; Beiki et al., 2012; Ebrahimi et al. 2011). The results showed that attribute subset selection can be beneficiary both to processing time and getting more accurate results. This reduces the dimensionality of the data and enables data mining algorithms to predict drought tolerance maize inbred lines faster and more effectively.

Attribute selection had an important effect on the classification and identification capability of drought tolerance inbred lines. Attribute weighting and selection methods based on SSR molecular marker data can classify drought tolerance and susceptible inbred lines (Table 2 and Table 3). The phi 033a3, bnlg1347a1, bnlg1347a2, bnlg172a2, bnlg381a3, bnlg381a2, bnlg172a3 and bnlg381a2 SSR fragments were the most important feature to distinguish drought tolerance from susceptible inbred lines, as defined by the entire attribute weighting algorithms (Table 2 and Table 3). These alleles can help to identify tolerance maize inbred lines from susceptible.

The goal of unsupervised pattern is to identify small subsets of alleles that display coordinated expression patterns. The unsupervised algorithm figure out the underlying similarities among a set of feature vectors, and to cluster similar vectors together (Ashrafi, Alemzadeh, Ebrahimi, Ebrahimie and Dadkhodaei 2011, Beiki, Saboor and Ebrahimi 2012, Garcia-Gonzalez et al. 2010), while supervised pattern like decision trees are very popular tools for classification (Migliorini et al. 2013, Yun et al. 2014). In unsupervised learning, the machine finds new patterns without relying on prior training examples, usually by using a set of pre-defined rules (Bravo and Medina 2008, El Harchaoui et al. 2013, Handfield et al. 2013, Markovich Gordon et al. 2012). Here we quantify the performance of a given unsupervised clustering algorithm applied to a given molecular marker data in terms of its ability to produce biologically meaningful clusters using a reference set of functional classes. We used three different unsupervised clustering methods (*K-Means*, *K-Medoids* and *SVC*) on 11 datasets created from SSR fragment allele attributes, which were assigned high weights. The performances of these algorithms varied significantly, usually these algorithms work well when the numbers of classes to be clustered are small. Here we have only two classes, tolerance and susceptible and it is expected that these algorithms are suitable for this condition and there is no need more complex clustering.

The results showed that combination of *K-Medoids* clustering method with Deviation attribute weighting was able to assign the drought tolerance inbred lines in right classes (Fig.2 -B) while Combination of *K-Means* clustering method with *PCA* attribute weighting selected the right classes of drought susceptible inbred lines (Fig.2 -D).

The main objective of decision analysis is to offer a theoretical representation of choices made in an environment of uncertainty (Berry and Linoff 2004, Senthil Kumar et al. 2013). The attractiveness of decision trees is due to the fact that, decision trees represent rules. Rules can readily be expressed so that humans can understand them (Berry and Linoff 2004). Decision trees provide the information about which attribute alleles are most important for prediction or classification. As shown in figure 3, 4 and 5, different algorithms of decision trees were able to produce the different decision trees with root and leaves. *Random Forest* model with *Accuracy* criteria on *Information Gain* and *Gini Index* datasets were able to produce decision trees with 100% overall accuracy and precision (Fig. 4). The most important attribute allele used to build this trees are bnlg1347a2 and bnlg381a3, which can be used to visually and explicitly represent decisions and decision making for susceptible and tolerance cultivars. Therefore, we are able to create a model that predicts the value of a target variable by learning simple decision rules inferred from the data features. It means simply by using 2 SSR markers, bnlg1347 and bnlg381, drought tolerance inbred lines can be predictable.

**Figure 5.**
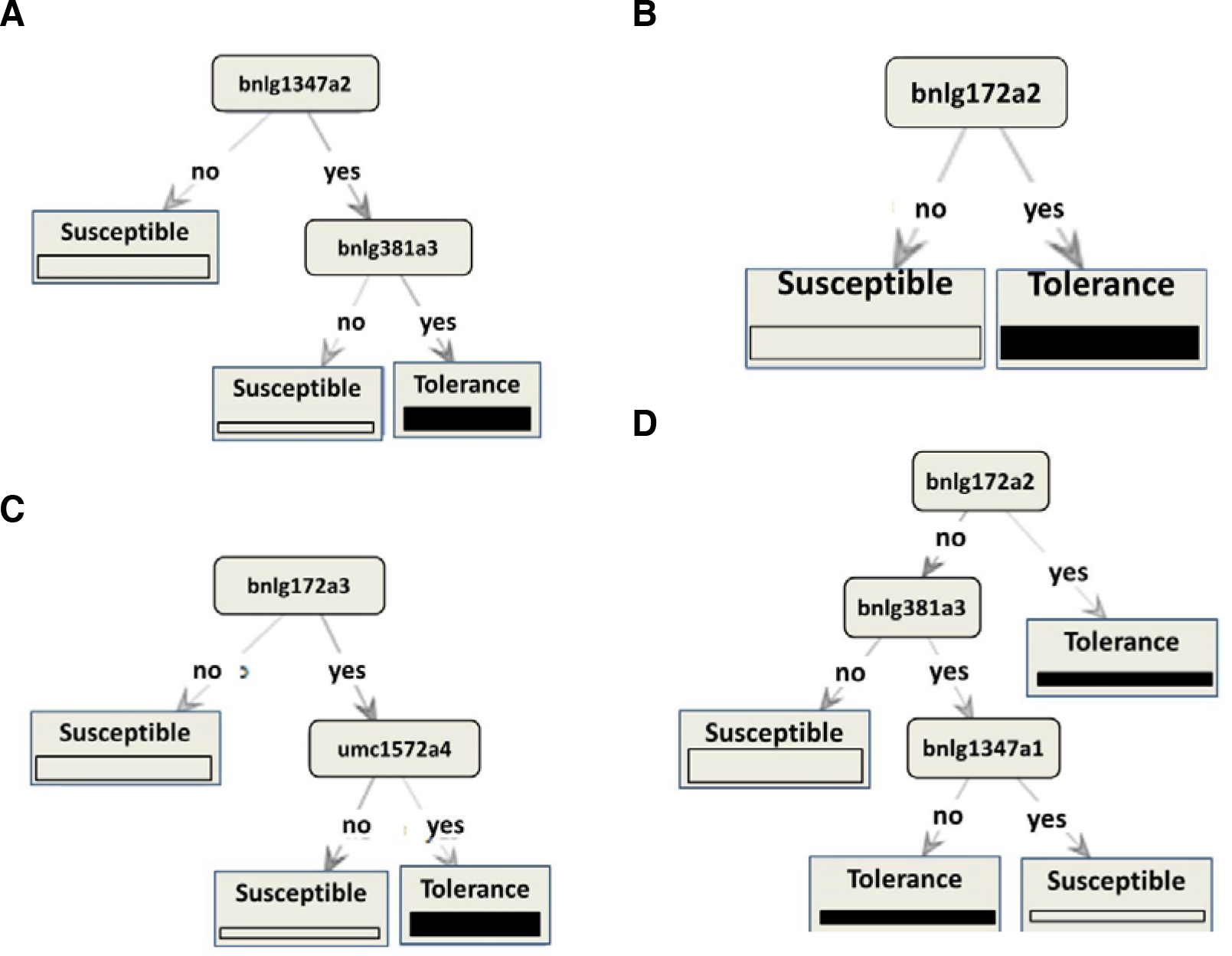
Random Forest model with Chi Square dataset when run on Accuracy (A), Gain Ratio (B) and Gini Index criteria (C); and with Information Gain dataset on Gini Index criterion (D).

Support Vector Machine are widely used in computational biology including genomics, proteomics, metabonomics due to their high accuracy, their ability to deal with high-dimensional datasets, and their flexibility in modeling diverse sources of data (Cai and Jiang 2013, O’Fallon et al. 2013, Verma and Melcher 2012, Xie et al. 2009). *SVM* established two-class classification-based Machine Learning methods can then be applied for developing an artificial intelligence system to classify a new allele or fragment into the member or non-member class. Our results suggested that only *SVM Evolution* algorithm with 100% overall accuracies and prediction (Table 6); would be the best candidate algorithm to predict drought tolerance inbred lines if they apply on *Chi Square* dataset.

*Naïve Bayes* based on *Bayes* conditional probability rule is used for performing classification tasks. *Naïve Bayes* assumes the predictors are statistically independent which makes it an effective classification tool that is easy to interpret (Paoin 2011, Prabhakara and Acharya 2012). Two models, *Naïve base* (returns classification model using estimated normal distributions) and *Naïve base kernel* (returns classification model using estimated kernel densities) used and the model accuracy in predicting the right resistance - susceptible computed as stated before.

The need to accelerate breeding for better adaptation to drought and other abiotic stresses is an issue of increasing urgency (Araus et al. 2008, Bänziger and Cooper 2001, Ribaut et al. 2009). Hence, as traditional breeding appears to be reaching a plateau; several approaches, which complement traditional with analytical selection methodologies, may be required to further improvement (Araus, Slafer, Royo and Serret 2008). The molecular approach has a great potential but actual results and delivery towards water limited environments are meager. Although the emergence new molecular techniques such as transcriptomics and proteomics propose a revolutionary impact in analytical breeding, DNA marker technology is still advantageous regarding cost/benefit and a potential partner for this recently introduced discipline. Inter-disciplinary scientists have been trying to understand and dissect the mechanisms of plant tolerance to drought stress using a variety of approaches. Application fields such as molecular genetics, combined with increasing computing power, supervised and unsupervised machine learning, can be used to identify groups of alleles with similar patterns of expression, and this can help provide answers to questions of how different alleles are affected by various traits and which alleles are responsible for specific hereditary characters. We are able to create models that have been applied successfully in the prediction, classification, estimation, and pattern recognition in abiotic stress. The molecular genetics road towards drought resistance is complex but we know that the destination is much simpler. One of the objective of this article was to address the need to bring to the molecular genetics community an increased understanding of knowledge discovery from data so that these robust computing paradigms may be used even more successfully in future molecular biology applications especially in abiotic and biotic stress arias.

## Acknowledgement

The author greatly appreciate support from department of Biology, Faculty of Science, University of Qom.

## Abbreviation

Fcdb: final cleaned database
SVM: Support Vector Machine
PCA: Principle Component Analysis
EM: Expectation Maximization

